# Cortical astrocytes control stress resilience

**DOI:** 10.64898/2026.01.17.700119

**Authors:** Veronika Kondev, Leanne M Holt, Brian Kipp, Trevonn M Gyles, Isla Racine, Jacob Abroon, Adam Ripp, William McKernan, Sumaiya Ahmed, Elizabeth Kahn, Angelica M Minier-Toribio, Clementine Blaschke, Sarah Naguib, Rita Futamura, Giselle Rojas, Alexa LaBanca, Eric J Nestler

## Abstract

**Background:** Chronic stress exposure is a risk factor for several psychiatric disorders, including post-traumatic stress disorder (PTSD) and major depression (MDD), with the prefrontal cortex (PFC) playing a key role in mediating this stress susceptibility. However, most individuals who are exposed to chronic stress are resilient and do not develop psychopathology. Recent evidence suggests that glial cells, especially astrocytes, play an important role in controlling stress-induced anxiety- and depression-like behavior, yet their role in contributing to stress resilience is not understood.

**Methods:** Using fiber photometry, chemogenetics, and RNA-sequencing in male mice, we establish a role for PFC astrocytes in stress resilience.

**Results:** We demonstrate that stress-induced increases in astrocytic calcium activity are both necessary and sufficient for resilience. Bioinformatic analysis reveals robust transcriptional responses in PFC astrocytes that differ between susceptible vs. resilient mice and are unique when compared to astrocytic transcriptional changes in other limbic regions. Comparison with human RNA-sequencing data indicates that molecular changes observed in PFC astrocytes from susceptible mice converge with gene expression changes observed in MDD patients.

**Conclusions:** Together, these data support targeting astrocytes as a potential therapy for negative behavioral consequences following stress exposure and reveal potential molecular mechanisms within PFC astrocytes that could contribute to depressive-like behaviors.

## Introduction

Stress is a major risk factor for a wide range of psychiatric disorders, including post-traumatic stress disorder (PTSD) and major depressive disorder (MDD)^1^. The prefrontal cortex (PFC) has been identified as a critical locus for controlling an individual’s responses to stress. Dysfunction within this region is strongly implicated in these stress-related conditions (reviews: ^2,3^). While the vast majority of studies have focused on neuronal populations in PFC and how they regulate stress effects, recent evidence suggests that glial cells, particularly astrocytes, also play an active role in the long-term behavioral and molecular consequences of stress exposure^4–7^.

A major function of astrocytes is their ability to monitor, integrate, and modulate neuronal activity, and this feature is heavily influenced by astrocytic calcium signaling^8–10^. Increased intracellular astrocytic Ca^2+^ is observed in rodent models of stress^11^, while human MDD and PTSD are associated with dysfunction in GFAP+ cells^5,12–15^. Furthermore, pharmacological or genetic blockade of astrocytic functions are sufficient to induce anhedonia-and anxiety-like behaviors, suggesting that astrocytes can causally modulate stress responses^16,17^. Indeed, studies have demonstrated that chemogenetic-induced increases in astrocytic Ca^2+^ reverse stress-induced deficits in sociability and anhedonia^18,19^, while decreasing astrocytic Ca^2+^ via overexpression of the Ca^2+^ pump PMCA can promote anxiogenesis^20^, suggesting bidirectional modulation of distinct symptoms related to MDD and PTSD. However, a role for astrocytes in modulating stress resilience is unknown.

Here, we causally link PFC astrocyte activity to stress resilience using the chronic social defeat stress (CSDS) model in mice^21^. Using in vivo fiber photometry and chemogenetic approaches, we reveal that stress-induced increases in astrocytic Ca^2+^ are necessary and sufficient to promote resilience. We further profiled transcriptional changes in sorted PFC astrocytes from susceptible vs resilient mice, and identified a stress resilience-specific transcriptional signature that distinguishes resilient mice from stress susceptible and stress naïve controls, such as processes related to serine-type peptidase activity, protein processing, and cytokine activity. Comparisons of our RNA-sequencing data to human MDD and PTSD datasets suggests that susceptibility-associated transcriptional changes in PFC astrocytes align with MDD, but not PTSD, gene patterns. Together, these findings support a role for stress-induced astrocytic transcriptional remodeling in MDD-related and propose new mechanisms for how astrocytes modulate cortical networks to control behavioral consequences of stress including the maintenance of a more resilient state.

## Materials and Methods

### Animals

All experiments were approved by Mount Sinai Institutional Animal Care and Use Committee, conducted in accordance with the National Institute of Health guidelines for the AALAC. Adult C57BL/6J wild-type male mice were used for all experiments. Mice were housed on a 12:12 hour light/dark cycle (07:00 lights on; 19:00 lights off) and were provided with food and water *ad libitum*. Mice that underwent surgery were 7-13 weeks old, and behavior was assessed at least 4 weeks following virus injection.

### Surgeries

Mice were anesthetized with an intraperitoneal injection of ketamine (100 mg/kg) and xylazine (10 mg/kg). PFC coordinates relative to bregma were: AP +2, ML ±0.7, DV −2.6 at a 10° angle.

### Viral Reagents

For fiber photometry experiments, mice were injected with AAV5-gfaABC1D-lck-GCaMP6f [GFAP-GCaMP] (Addgene) in the PFC. To inhibit astrocytic Ca^2+^ levels from increasing, AAV5-GfaABC1D-mCherry-hPMCA2w/b [PMCA] was injected. For chemogenetic activation of astrocytes, AAV5-GFAP-hM3D(Gq)-mCherry or control (AAV5-GFAP-mCherry) was injected into the PFC.

### Fiber Photometry Recordings and Analysis

For cell body recordings of PFC astrocyte activity, the fiber photometry system (FP3002 system from Neurophotometrics, NPM) was used. Analysis was performed as previously described^22^.

### Chronic Social Defeat Stress (CSDS) and Social Interaction Test (SIT)

C57 mice were introduced into the cage of resident CD1 retired breeder mice for 5 min/day over 10 days^7,21^. The SIT for social avoidance was performed within 24 hours after the last CSDS session. Social interaction ratio was calculated by dividing the time spent in the interaction zone when the target mouse was present divided by the time spent in the zone when the target mouse was absent. Mice with a SI ratio < 0.9 were susceptible, and with > 1.1 were resilient^7,21^.

### Astrocyte Isolation

Astrocytes (n = 7–8/group) were isolated using Miltenyi BioTech’s ACSA-2 MicroBead Kit as previously described ^23,24^ from freshly micropunched (unilateral, 12G) PFC tissue^7,23,25^.

### RNA extraction, Library preparation, and Sequencing

RNA was extracted as previously reported^7^ using Zymo’s Directzol RNA MicroPrep kit following manufacturer’s instructions. RNA quality and quantity were assessed using Bioanalyzer (Agilent). 500 pg of RNA was used as input in Takara’s SMARTer® Stranded Total RNA-Seq Kit v2 – Pico, with ribodepletion. Sequencing libraries were generated for each sample individually using Takara’s Unique Dual Index Kit. Following library preparation, sequencing was performed with Azenta on an Illumina Novaseq. Differential gene expression (DEG) was performed in R using the DESeq2 package. Significant DEGs were determined by 20% log2foldchange and p < 0.05. Volcano plots, heatmaps, Venn diagrams, and gene ontology plots were generated using tidyverse and ggplot. Rank rank hypergeometric overlap (RRHO) were created using RRHO2 package. Gene ontology (GO) was performed using enrichR package. Top 10 terms were graphed presented and summarized for selected GO terms. Full GO terms are in supplemental figures. For comparisons with human MDD and PTSD data sets, data was taken from Girgenti, et al.^26^ Supplemental Table. For correlations with behavior, variance-stabilized counts (VST) were obtained from DESeq2 analysis. For each gene, Pearson correlation coefficient was computed between its VST-normalized expression values and the corresponding behavioral metric. Correlations were calculated using pairwise complete observations to handle missing values.

### Statistical Analyses

All data are plotted as mean +/- SEM, and statistical analysis was performed in Prism 10.

## Results

### Astrocytes in PFC respond to stress exposure

To first examine how cortical astrocytes respond during stress, we injected an astrocyte-specific calcium indicator, AAV5-gfaABC1D-lck-GCaMP6f [GFAP-GCaMP], into the PFC and used fiber photometry to record changes in astrocytic Ca^2+^ levels during CSDS (Fig. 1A-B). During daily 5 minute physical attacks by a CD1 aggressor, we observed significant increases in PFC astrocyte Ca^2+^ levels (Fig. 1C-D). A separate cohort of mice were co-injected with GFAP-GCaMP and the calcium extruder, AAV5-gfaABC1D-mCherry-hPMCA2w/b [PMCA], to prevent increases in intracellular Ca^2+^ levels. PMCA significantly blunted attack-evoked Ca^2+^ increases, validating that the stress-induced Ca^2+^ signals observed are not simply the result of fiber optic cable movement artifacts (Fig. 1C-D). This was specific to attacks, as non-attack interactions (i.e., sniffing, grooming, etc.) did not elicit changes in astrocytic Ca^2+^ signaling (Fig. 1E).

**Figure 1:**
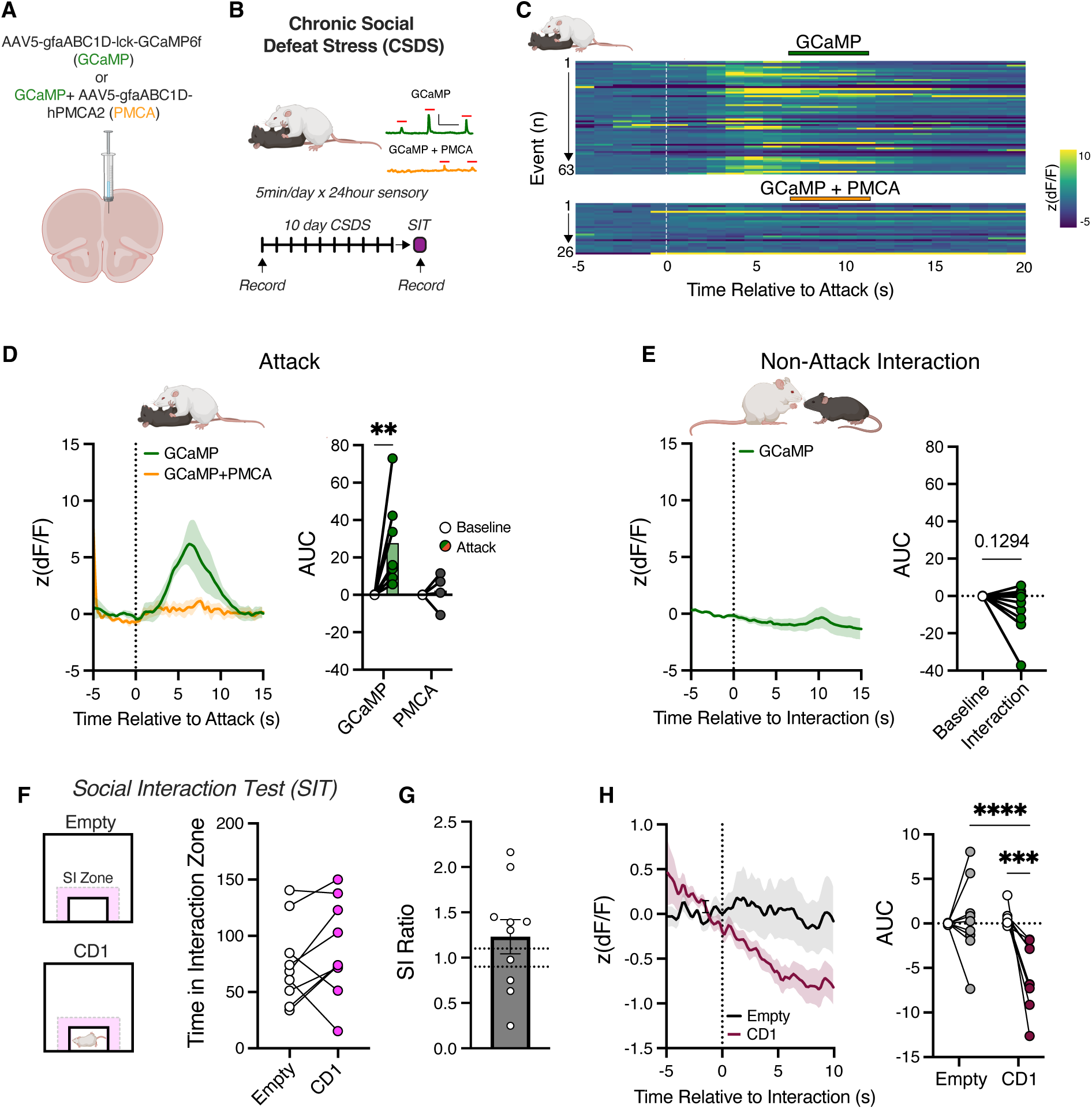
PFC astrocytes respond differently to social defeat stress vs a social interaction test after stress exposure. A) Schematic for experimental design of recordings of Ca^2+^ from PFC astrocytes with or without PMCA. B) Chronic social defeat stress (CSDS) paradigm and when fiber photometry recordings were conducted. C) Heatmap of change in astrocyte Ca^2+^ activity in response to attacks during each social defeat exposure. D) Average change in astrocyte activity in response to attack (left) and resulting area under the curve (AUC) following attack. 2-Way ANOVA, F_Attack_(1,9)=5.46; p=0.0443. E) Average change in astrocyte activity during non-attack interactions during day 1 of CSDS and resulting AUC. Paired t-test, p=0.1294. F) Social interaction test (SIT) schematic and time spent in social interaction zone in fiber photometry mice. G) Resulting social interaction (SI) ratio. H) Change in astrocyte activity during entry to the interaction zone (SI Zone) when the chamber is empty vs when a CD1 mouse is present and resulting AUC. 2-Way ANOVA, F_Interaction_(1,16)=11.43; p=0.0038.

We confirmed that stress exposure, in general, enhances PFC astrocyte Ca^2+^ levels. First, we subjected mice to a single session of fear conditioning, during which they were exposed to six tone-shock pairings (Fig. S1A). We observed that footshock, but not the tone, significantly increased astrocyte Ca^2+^ levels, and this effect too was blunted by PMCA co-expression (Fig. S1B-D). We also observed significant increases in astrocyte Ca^2+^ activity in “anxiogenic” environments in the elevated plus maze (EPM). When mice entered the open arm, there was a significant increase in astrocytic Ca^2+^ levels compared to when mice entered the closed arm (Fig. S1E). Together, these data demonstrate that astrocytes in the PFC are activated by stress and anxiogenic stimuli.

### Distinct changes in PFC astrocyte activity during social interaction testing

We next measured PFC astrocyte activity during a social interaction test after ten days of CSDS, which is used to behaviorally phenotype mice into stress susceptible vs stress resilient groups^21^ (Fig. 1F). Mice that underwent CSDS were first exposed to an open field chamber with no target mouse present in the social interaction grid (empty); and then immediately exposed to the same arena with a novel CD1 mouse contained within the interaction grid (CD1). We observed a decrease in astrocyte Ca^2+^ activity in PFC upon the test mouse’s entry into the interaction zone when interacting with a CD1 mouse compared to interactions with the empty chamber (Fig. 1G-H). However, the changes observed in astrocyte signaling during the social interaction did not reflect phenotypic profiling of resilience vs. susceptibility (Fig. S2A-C). Together, these data demonstrate that stressful stimuli, but not their cues, enhance PFC astrocyte Ca^2+^ activity, and that astrocyte signals during the social interaction test do not differ between susceptible and resilient mice.

### Chemogenetic activation of PFC astrocytes promotes stress resilience

Given that stress exposure enhances astrocytic Ca^2+^ levels only during the stressor, we causally test the role of this response in modulating stress susceptibility vs resilience. First, we used chemogenetics to pharmacologically elevate astrocytic Ca^2+^ levels during the CSDS procedure. Mice were virally injected with the excitatory Gq-DREADD [AAV5-GFAP-hM3D(Gq)-mCherry] or control [AAV5-GFAP-mCherry] (Fig. 2A); following virus expression, mice were exposed to ten days of CSDS during which mice were injected (*i.p.*) with deschloroclozapine (DCZ) 15 min before each social defeat session (Fig. 2B). 24 hours after the last day of CSDS (day 10), mice were given a social interaction test (day 11). We found that increasing astrocytic Ca^2+^ levels during stress increased social interaction on the test day (Fig. 2C). This was reflected in a significant increase in the proportion of resilient mice, demonstrating that astrocytic Ca^2+^ signaling during stress serves to promote stress resilience and counteract negative behavioral sequelae (Fig. 2D).

**Figure 2:**
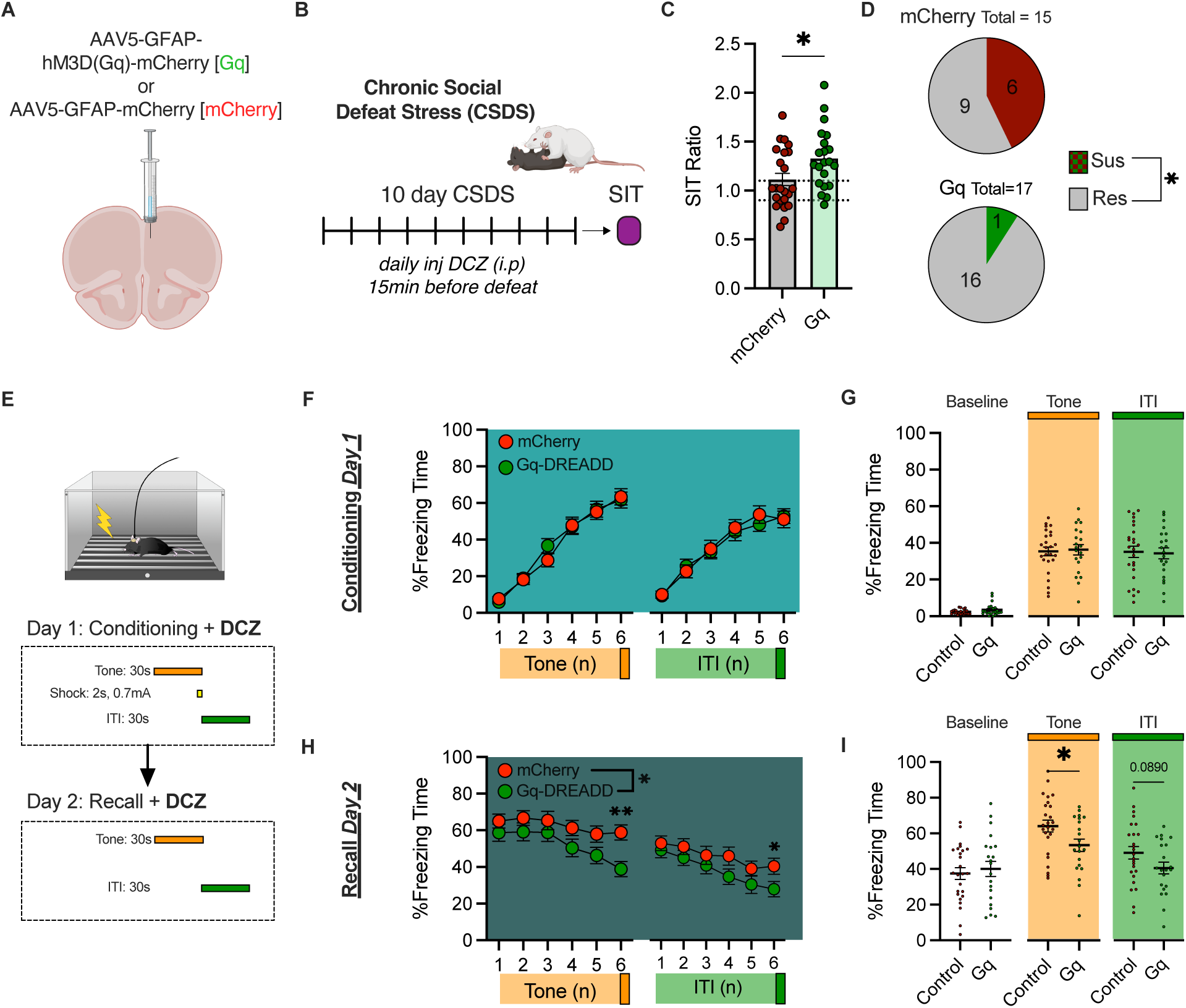
Chemogenetic activation of PFC astrocytes promotes stress resilience. A) Schematic of viral design. B) Schematic of CSDS protocol and social interaction test (SIT). C) SIT ratio following chemogenetic activation of astrocytes during defeat (unpaired t-test; t=2.317, df=42; p=0.0254). D) Proportion of stress susceptible vs. resilient mice (Chi-square; 5.428, 1; p=0.0198). E) Fear conditioning and recall protocol. F) % time spent freezing over tone and intertrial interval (ITI) trials. G) Average % time freezing during baseline (pre-shock), tone, and ITI. H) % time freezing over tone and ITI trials during recall test (2-Way ANOVA; F_Virus_(1,44)=4.462; p=0.0404). I) Average % time freezing over different stimuli; unpaired t-test (tone, t=2.240, df=44; p=0.0302).

To test whether this effect was specific to behavioral adaptations to stress or rather a general influence on learning and memory, we exposed the same mice to fear conditioning and recall (Fig. 2E). On day 1, mice were injected with DCZ and exposed to six tone-shock pairings. We did not observe any significant differences in baseline freezing to the new context or the conditioned stimulus (tone), or in generalized freezing during the intertrial interval (ITI), compared to controls (Fig. 2F-G). The next day, mice were injected with DCZ and placed back in the same context to assess recall of both contextual and conditioned freezing behavior. Astrocytic Gq-DREADD activation promoted extinction of tone-elicited freezing within the session, resulting in an overall decrease in freezing time during the tone (Fig. 2H-I). These data suggest that astrocytic Ca^2+^ levels could drive resilience by disrupting the encoding of stress-cue associations or facilitating extinction of fear memories that drive social avoidance after CSDS.

### Inhibiting PFC astrocytic Ca^2+^ signaling promotes susceptibility

We next tested whether astrocytic Ca^2+^ levels can bidirectionally control resilience by inhibiting astrocytic Ca^2+^ and testing whether that would reduce resilience, i.e., drive susceptibility. We injected mice bilaterally with viral vectors that express PMCA in astrocytes, which we have shown reduces intracellular Ca^2+^ mobilization in PFC astrocytes (Fig. 1). Following viral expression, mice were exposed to ten days of CSDS and a subsequent social interaction test (Fig. 3A-B). PMCA mice exhibited significantly lower social interaction scores and trends towards increased susceptibility (Fig. 3C-D), confirming bidirectional modulation of stress adaptation following astrocytic Ca^2+^ disruption: increasing Ca^2+^ with Gq-DREADDS promotes resilience (Fig. 2), while inhibiting Ca^2+^ signaling biases towards susceptibility (Fig. 3).

**Figure 3:**
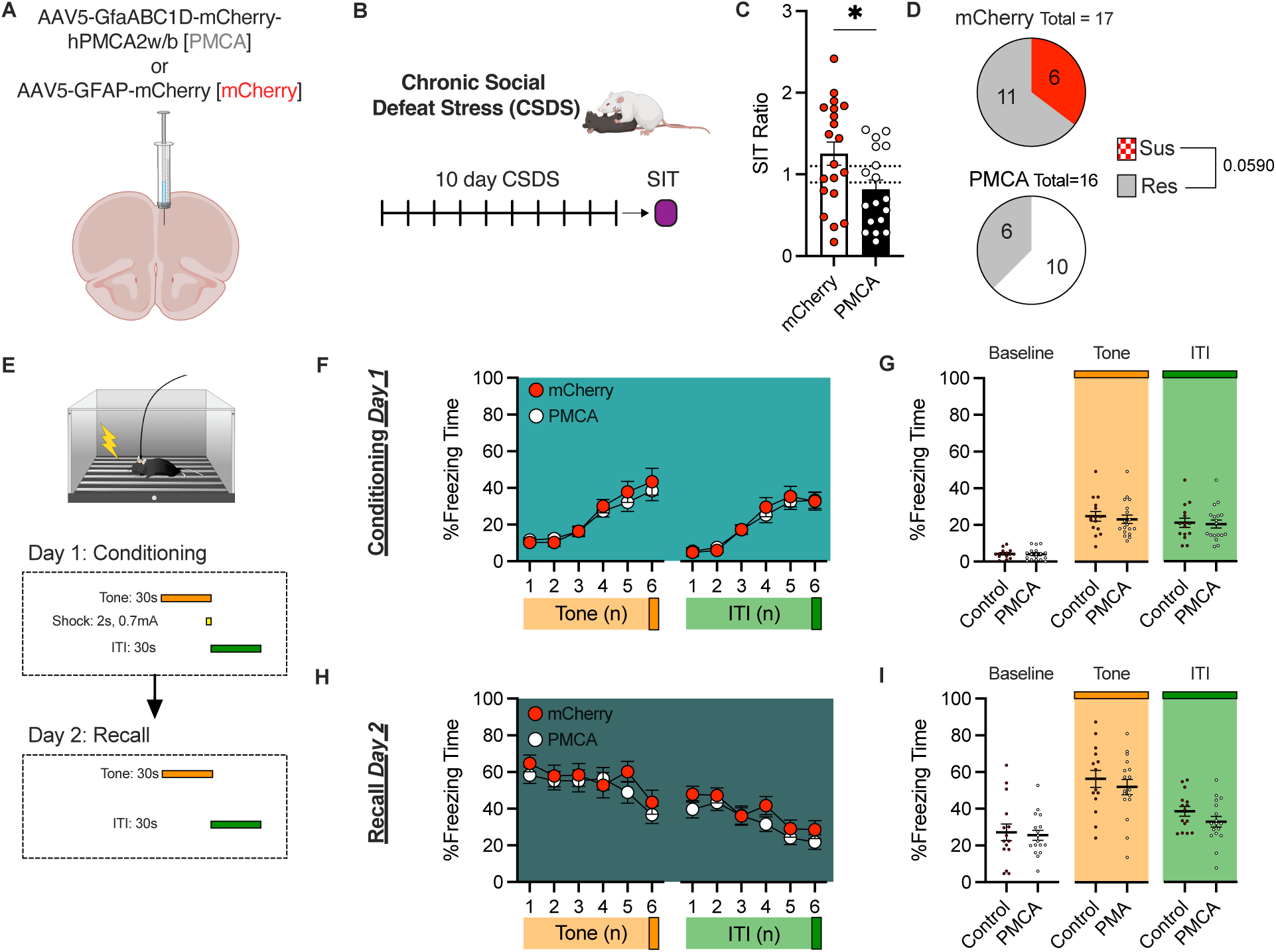
Astrocytic disruption of calcium signaling reduces social interaction with no effect on fear conditioning or recall. A) Schematic of viral vectors injected into PFC to reduce astrocytic Ca^2+^ signaling. B) CSDS paradigm. C) SIT ratio (unpaired t-test; t=2.359, df=36; p=0.0239). D) Proportion of stress susceptible vs. resilient mice (Chi-square, 2.443, 1; p = 0.0590). E) Fear conditioning and recall protocol. F) % Time freezing during tone and ITI trials. G) Average % time freezing during conditioning. H) %Time freezing during tone and ITI trials during recall. I) Average % time freezing during recall.

We additionally investigated general learning and memory deficits using fear conditioning, as described before. Surprisingly, PMCA had no significant effect on baseline freezing, conditioned freezing, or generalized freezing across either conditioning or recall (Fig. 3E-I), suggesting that tonic inhibition of PFC astrocytic Ca^2+^ signaling does not shift acute fear memory acquisition or expression, but does impair behavioral adaptation following chronic stress.

### Dramatic regulation of the PFC astrocyte transcriptome in stress resilient mice

Astrocyte Ca^2+^ signaling is linked to many downstream functions, including changes in gene expression and subsequent modulation of neuronal activity^8–10^. Therefore, we explored molecular correlates of stress-induced adaptations in PFC astrocytic Ca^2+^ signaling. We carried out bulk RNA-sequencing of sorted PFC astrocytes after CSDS to characterize transcriptional regulation in stress resilience vs susceptibility. Wild-type male mice were exposed to 10 days of CSDS and subsequent a subsequent social interaction test, and 24-48 hours later PFC tissue was isolated and astrocytes were sorted using MACs (Fig. 4A-C); nucleus accumbens (NAc) punches were simultaneously processed from the same mice and previously published^7^. We observed significant transcriptional changes in PFC astrocytes from stress susceptible vs stress resilient mice, with the largest number of differentially expressed genes (DEGs) found in resilient mice compared to unstressed, control mice (Fig. 4D-H). By contrast, we observed one fifth fewer DEGs in susceptible mice compared to controls, with minimal overlap seen between resilient and susceptible DEGs (Fig. 4G).

**Figure 4:**
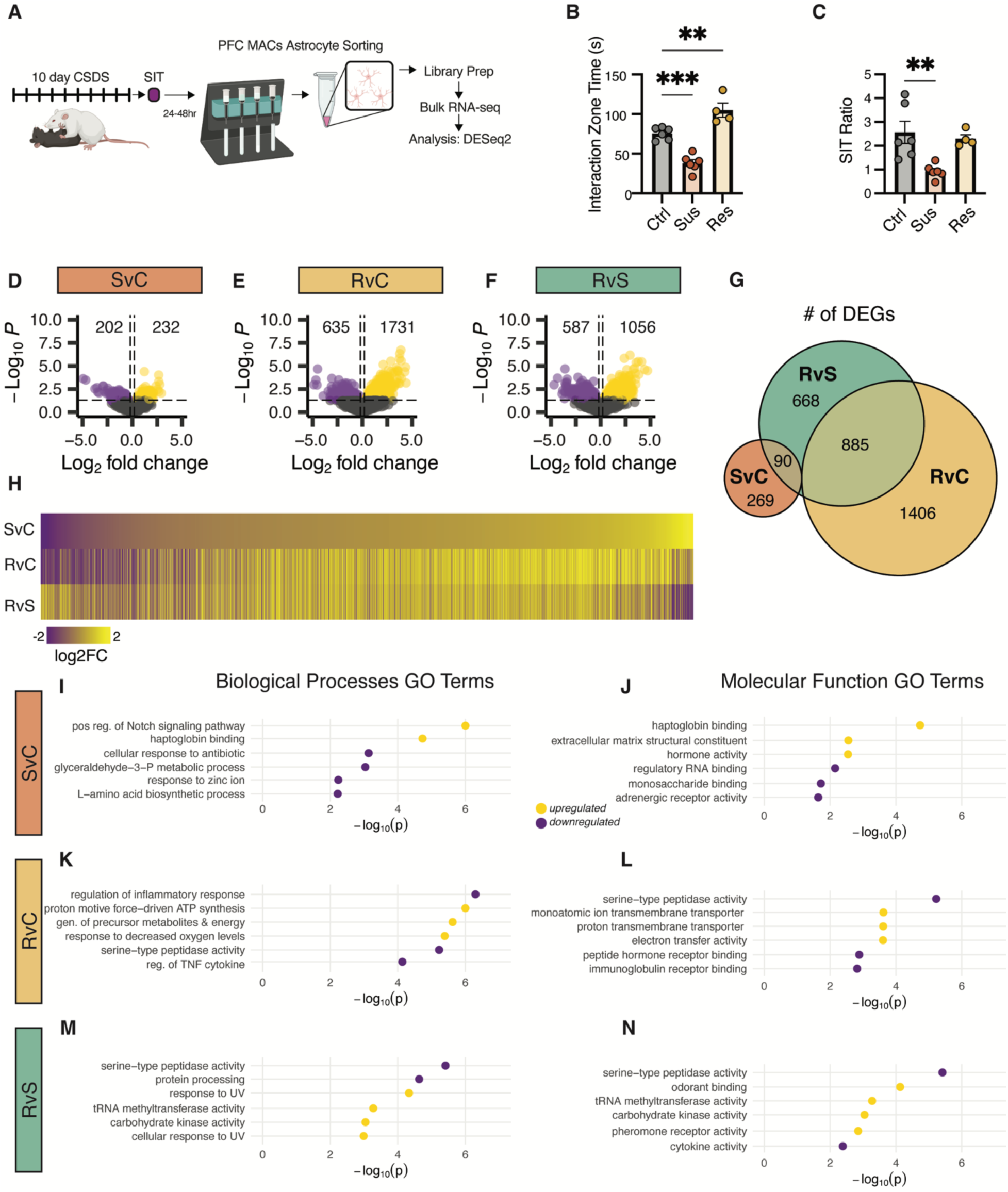
Transcriptomic changes induced in PFC astrocytes of stress resilient vs susceptible mice. A) Schematic of MACs and RNA-sequencing of PFC astrocytes. B) Interaction zone time from sequenced mice (One-Way ANOVA, F(2,13)=39.51, p<0.0001). C) Social interaction ratio (SIT ratio) (One-Way ANOVA, F(2,13)=7.965; p=0.0055). D) Volcano plot of differentially expressed genes (DEGs) comparing susceptible vs. control mice. E) Volcano plot of DEGs comparing resilient vs. control. F) Volcano plot from resilient vs. susceptible mice. G) Overlap of DEGs across comparisons. H) Heatmap of changes in DEGs using susceptible vs. control as reference. I) Selected gene ontology (GO) biological processes terms of DEGs from susceptible vs. control comparisons. J) Selected GO molecular functions from susceptible vs. control mice. K) Biological processes GO terms from resilient vs. control. L) Molecular function GO terms from resilient vs. control. M) Biological processes GO terms from resilient vs. susceptible. N) Molecular function GO terms from resilient vs. susceptible.

Comparison of gene ontology (GO) terms further confirms that DEGs induced in PFC astrocytes of susceptible and resilient mice associate with distinct downstream biological processes and molecular pathways (Fig. S3). For example, susceptibility is associated with developmental, structural, and hormonal changes, while resilience is associated with downregulation of genes related to *inflammatory responses* when comparing to unstressed, control mice (Fig. 4I-L). Directly comparing resilient to susceptible mice reveals that resilience promotes a downregulation in *serine-type peptidase activity*, *cytokine signaling*, and *protein processing* (Fig. 5M,N).

**Figure 5:**
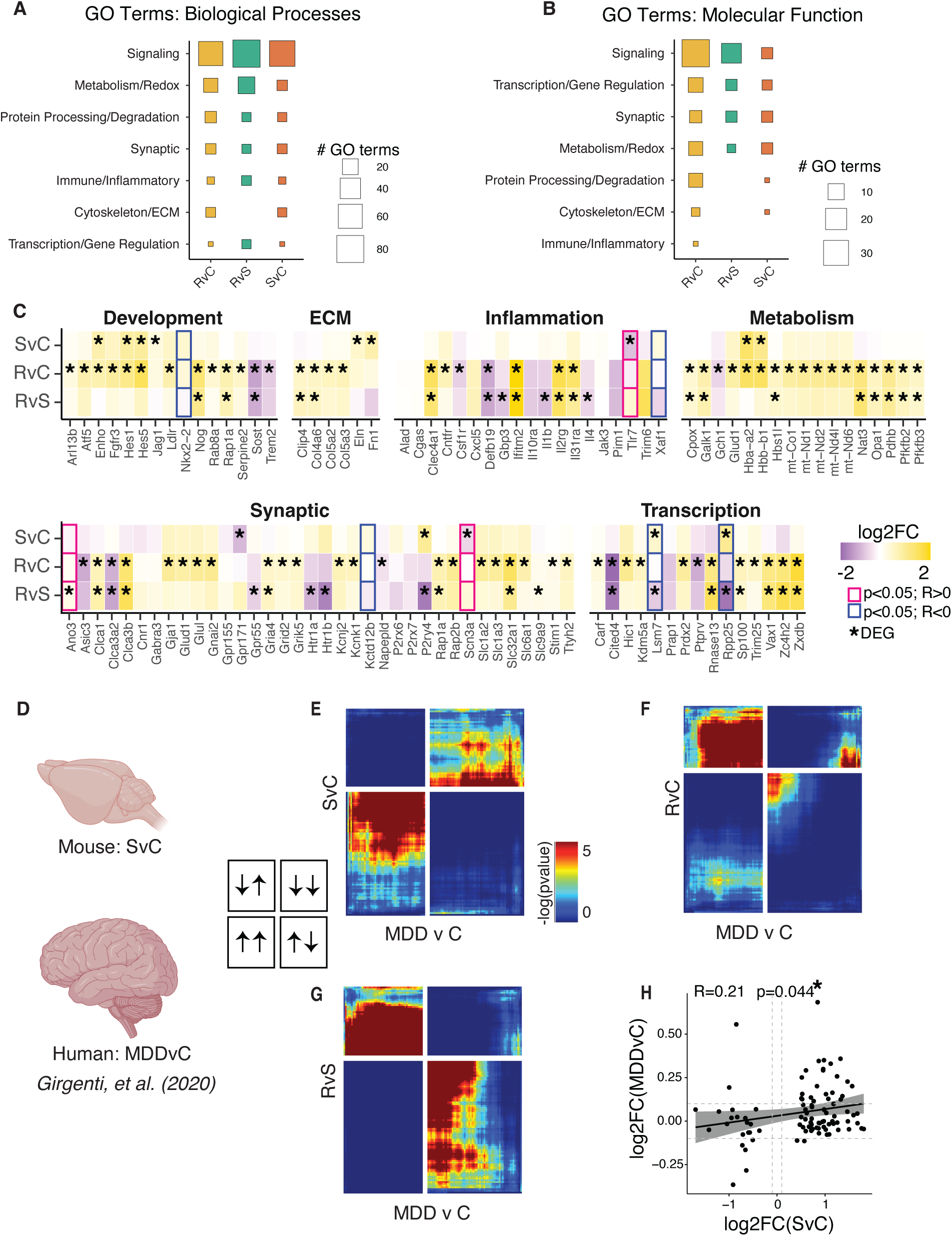
Transcriptional changes in PFC astrocytes of susceptible mice show convergence with human MDD, but not PTSD. A) Number of biological processes GO terms associated with categories across comparisons. B) Number of molecular function GO terms associated with categories across comparisons. C) Heatmap of log2foldchange of key genes associated with biological functions; outline indicates whether gene significantly correlates with social interaction ratio, while * indicates significant DEG. D) PFC susceptible vs control (SvC) gene expression was compared with a previously published human dataset assessing gene changes in dorsolateral PFC of humans with major depression disorder (MDD) vs healthy controls (Girgenti et al., 2020). E) RRHO2 plot comparing SvC and MDDvControl. F) RRHO2 plot comparing resilient vs control (RvC) and MDDvControl. G) RRHO2 plot comparing RvS and MDDvControl. H) Correlation of log2foldchange of significant DEGs from SvC comparisons and MDDvC.

The number of DEGs shared across resilience reveals large overlap (885 DEGs) whether comparing resilient mice to unstressed controls or to susceptible mice (Fig. 4G). GO analysis of these conserved resilient genes reveals shared biological and molecular mechanisms related to *metabolism*, *apoptosis*, and *chloride channel activity* (Fig. S4A-D). We also found conserved stress responses that include *Notch* signaling and *redox* function (Fig. S4E-F). Together, these reveal potential biological processes and functions within PFC astrocytes that could underlie stress resilience.

We further assessed genes related to susceptibility or resilience across distinct molecular and biological functions and correlated changes in gene expression with social interaction ratios (Fig. 5A-C). We identified several key genes that correlate with behavior which could alter astrocyte function and astrocyte∼neuron interactions. For example, *Tlr7,* which positively correlated with social interaction, is an innate immune receptor whose activation shapes astrocytic inflammatory responses^27^. Increased *Tlr7* expression in PFC astrocytes of resilient mice may reflect an astrocytic state that is better equipped to modulate or constrain stress-induced inflammatory signaling. We also identified *Ano3*, a member of the anoctamin (TMEM16) family linked to Ca² -dependent chloride conductance. *Ano3* levels positively correlated with social interaction ratios and were significantly upregulated in resilient animals, suggesting a potential shift in astrocytic Ca² -responsive signaling states that may support improved neuron–glia communication under stress. Together, these findings highlight genes in PFC astrocytes that track with individual differences in behavioral susceptibility vs resilience and may contribute to mechanisms that underlie stress resilience.

### Stress susceptibility and resilience associate with distinct molecular changes across limbic brain regions

We next compared DEGs induced in the PFC with previously published data taken from the same mice assessing transcriptomic changes in NAc astrocytes^7^ (Fig. S5). Generally, there were more DEGs induced in the NAc than in the PFC, with similar proportions of up- and down-regulated genes across regions (Fig. S5A-C). We observed very little overlap in DEGs between the PFC and NAc across conditions, demonstrating that stress susceptibility and stress resilience induce mostly distinct transcriptomic changes in different brain regions (Fig. S5D-F). Rank rank hypergeometric overlap (RRHO) confirms discordance in gene patterns across the PFC vs NAc (Fig. S5G); we found the most discordance in resilient vs. control mice demonstrating that stress adaptations leading to resilience are highly distinct across brain regions (Fig. S5H-I).

### Molecular changes in susceptible but not resilient mice converge with human MDD-associated gene expression abnormalities

Finally, we wanted to assess whether stress-induced changes in PFC astrocytic gene expression correlate with human pathology. While stress has been linked to anhedonia, anxiogenesis, and generalized or irrational fear responses, these core symptoms represent distinct symptom classes of psychiatric disorders. Low mood, anhedonia, and suicidality are core symptoms in MDD; while hyperarousal, vigilance, and trauma-related reactivity are more characteristic of PTSD. To assess whether molecular changes induced in PFC astrocytes of mice exposed to CSDS correlate with such distinct stress-induced sequalae in humans, we compared PFC astrocyte transcriptome patterns from susceptible or from resilient mice (vs. controls) with a previously published dataset from the dorsolateral PFC (dlPFC) (vs. healthy controls) of MDD or PTSD male patients^26^ (Fig. 5D).

We observed significant concordance in transcriptomic profiles of PFC astrocytes from susceptible mice compared to human MDD, demonstrating that gene expression changes seen in susceptible mice parallel those observed in human MDD (Fig. 5E). On the other hand, when comparing MDD with resilient vs control mice or resilient vs susceptible mice, we found robust discordance, further supporting the view that transcriptomic changes in PFC astrocytes from susceptible mice may reflect conserved vulnerability mechanisms across species (Fig. 5F-G). Indeed, when correlating directly for changes in gene expression, we found significant positive correlation between DEGs in PFC astrocytes of susceptible mice and in bulk PFC of human MDD patients (Fig. 5H).

We repeated this analysis to compare molecular changes in PTSD patients (vs healthy controls) and observed very different results (Fig. S6A-B). Transcriptomic changes observed in PFC astrocytes of susceptible mice did not significantly correlate with PTSD patients. Together, this suggests that the astrocytic molecular landscape induced by CSDS aligns more closely with MDD-related dysfunction, supporting the idea that astrocyte-mediated impairments may be core features of depression vulnerability, rather than generalized features of trauma exposure or PTSD-specific domains.

## Discussion

Astrocytes are known to modulate baseline emotional behaviors, as well as anxiety- and depression-like behavioral sequelae after stress. Here, using the CSDS model, we refine the role for Ca^2+^ signaling in PFC astrocytes in controlling stress susceptibility vs resilience in male mice and define the transcriptomic profiles associated with these distinct responses to chronic stress. We also reveal that molecular changes in PFC astrocytes from susceptible mice resemble changes observed in MDD patients, demonstrating the human relevance of our findings in mice.

First, we show that stressful and anxiogenic stimuli promote increases in PFC astrocytic Ca^2+^ signaling. Our fiber photometry data, wherein we observed increases in astrocytic Ca^2+^ during stress exposures, but not the social interaction test after stress, suggest that stress-induced Ca^2+^ activity serves to constrain the negative effects of stress and prevent social avoidance. It is noteworthy that our fiber photometry data demonstrate a decrease in astrocyte Ca^2+^ activity during the social interaction test upon entry to the interaction zone when a CD1 aggressor is present, an effect that was equivalent in susceptible and resilient mice (Fig. 1). PFC astrocyte activity is increased during social interaction in control mice, but corticosterone treatment abolishes this effect^18^. This suggests that the lack of differences across susceptible vs. resilient mice during the social interaction test could be due to a general effect of stress-induced elevations in corticosterone signaling after CSDS. Indeed, it has been shown that susceptible and resilient mice do not significantly differ in their corticosterone levels after stress^21,28^. Previous work also demonstrates that astrocytes respond to several stressors, including social defeat and footshock^29^ as we show here, as well as anxiogenic environments like the open arm of the elevated plus and zero mazes^20^.

Our astrocyte manipulation data demonstrate that increases in PFC astrocytic Ca^2+^ levels are both necessary and sufficient for stress resilience. Chemogenetic activation of astrocytes across ten days of CSDS increases social interaction and resilience, whereas reduction in tonic astrocyte Ca^2+^ levels via PMCA overexpression reduces social interaction and resilience. It has been demonstrated previously that chemogenetic activation of PFC astrocytes has no effect on basal anxiety-like behavior or social avoidance^20^, while others have shown that such activation is anxiolytic but pro-depressive, including reduced social interaction^18^. Our data indicate that chronic stress may reprogram PFC astrocytes so that activation shifts from pro-depressive at baseline, to anti-depressive following stress, in parallel with previous studies reporting anti-depressive effects after corticosterone treatment^18^. However, how this shift occurs, as well how astrocytes shape cortical neural activity, remains to be determined.

Our RNA-sequencing data suggest several distinct biological and molecular pathways that might underlie stress susceptibility vs stress resilience, such as inflammation, structural changes, and oxidative processes, among many others. Future studies will validate causal roles for individual genes in these pathways, as well as how cortical networks are dysregulated as a result. Our fear conditioning data, demonstrating that chemogenetic activation of PFC astrocytes is sufficient to decrease conditioned fear during recall, suggest that astrocytic Ca^2+^ signaling could generally disrupt fear learning and memory processes. However, whether this is due to an encoding or extinction deficit remains to be determined.

Finally, our RNA-sequencing reveals several further conclusions. First, CSDS induces many-fold more gene expression changes in PFC astrocytes in resilient mice vs susceptible mice. Greater transcriptional responses in stress resilience have been observed in PFC and in several other limbic brain regions by bulk RNA-sequencing, which includes numerous neuronal-enriched genes as well^21,30,31^. Thus, the conclusion that stress resilience represents the more plastic state and that stress susceptibility may reflect a failure of this plasticity holds across neurons and glia. Second, molecular changes induced in PFC astrocytes by susceptibility or by resilience are divergent from those observed in NAc astrocytes of the same mice, suggesting clear limbic-dependent transcriptional changes. This observation is in line with many studies demonstrating region- and circuit-dependent synaptic changes on a neuronal level and also suggests similar regional specificity for astrocytes. However, the exact relationship between astrocytes and neurons, and how they influence each other to elicit such behavioral effects, remains to be determined. Third, the gene expression changes induced in PFC astrocytes in susceptible mice, but not those observed in resilience mice, resemble changes observed in bulk RNA-sequencing of the PFC of MDD patients. This finding adds to previous studies which also showed strong concordance between bulk transcriptomic abnormalities in limbic brain regions of chronically-stressed mice and the homologous regions from MDD patients^32,33^ and demonstrates that part of this convergence occurs in astrocytes. Such convergence further establishes that chronic stress in mice induces a significant portion of the molecular pathology seen in human depression.

The observed convergence in gene expression abnormalities in PFC astrocytes of susceptible mice and in PFC of humans with depression but not with PTSD suggests that the molecular adaptations we observe here could underlie susceptibility for specific stress-induced symptoms such as anhedonia that reflect MDD. One important limitation of our current work is the inclusion of only male mice. Future studies will need to determine the role of PFC astrocytes in modulating stress resilience vs susceptibility, as well as sex-specific transcriptional responses in astrocytes that regulate responses to chronic stress.

## Conclusions

Here we identify a role for astrocytic Ca^2+^ signaling in mediating stress resilience. Increases in astrocytic Ca^2+^ levels are necessary and sufficient to promote resilience after chronic stress, as well as decrease conditioned freezing, suggesting dysfunctions in learning and memory mechanisms that drive social avoidance. We demonstrate potential molecular mechanisms, that converge with human MDD patients, that could drive astrocytic involvement in resilience. Together, these data demonstrate that astrocytes are key players in adaptive vs maladaptive behavioral sequelae after chronic stress and highlight key processes underlying MDD that could be targeted for therapeutic purposes.

## Acknowledgements

This work was supported by funding from the NIH (R01MH129306) and the Hope for Depression Research Foundation to E.J.N, as well as T32 grant support (5T32MH13583-02) to VK. The authors would like to thank Kinnerett Rosen and Ezekiell Mouzon for animal husbandry.

## Data availability

All RNA-seq data reported in this study will be deposited in the Gene Expression Omnibus (GEO).

