## Supplemental Methods for "Cortical astrocytes control stress resilience"

**Supplementary Methods**

**Animals**

Adult C57BL/6J wild-type mice were used for all experiments. Mice were housed on a 12:12 hour light/dark cycle (07:00 lights on; 19:00 lights off) and were provided with food and water *ad libitum*. Mice that underwent surgery were 7-13 weeks old, and behavior was assessed at least 4 weeks following virus injection. All experiments were approved by Mount Sinai Institutional Animal Care and Use Committee, conducted in accordance with the National Institute of Health guidelines for the AALAC.

**Surgeries**

Mice were anesthetized with an intraperitoneal injection of ketamine (100 mg/kg) and xylazine (10 mg/kg), then head-fixed in a stereotaxic apparatus (Kopf Instruments). Syringe needles (33G, Hamilton) were used to infuse 1 μl of virus at a 0.1 μl/min flow rate unilaterally for fiber photometry experiments, or bilaterally for all other experiments. Needles were kept in place for 10 minutes after injection before being retracted to allow for virus diffusion. Coordinates for PFC were as follows, from bregma: 10° angle, AP +2, ML ±0.7, DV -2.6mm. For fiber photometry, 400 μm-wide optical fibers (Doric, MFC_400/430- 650 0.66_4.5 mm_MF2.5_FLT) were unilaterally implanted at AP +2, ML ±0.7, DV -2.6mm, 0° angle. Optical fibers were secured in place using dental cement (3M) and covered with a layer of black dental cement (C&B Metabond). Virus infusion and optical fiber placement was confirmed either by immunohistochemistry on fixed brains sections or by dissection of fresh tissue under fluorescent light.

**Viral Reagents**

For fiber photometry experiments, mice were injected with AAV5-gfaABC1D-lck-GCaMP6f [GFAP-GCaMP] (Addgene) in the PFC. To inhibit astrocytic Ca^2+^ levels from increasing, AAV5-GfaABC1D-mCherry-hPMCA2w/b [PMCA] was injected. For chemogenetic activation of astrocytes, AAV5-GFAP-hM3D(Gq)-mCherry or control (AAV5-GFAP-mCherry) was injected into the PFC.

**Fiber Photometry Recordings and Analysis**

For cell body recordings of PFC astrocyte activity, the fiber photometry system (FP3002 system from Neurophotometrics, NPM) was used. Low-autofluorescent patch-cords (Doric, MFP_400/430/1100-0.66_3m_FCM-MF2.5_LAF) were used according to manufacturer’s instructions with the FP3002 Bonsai node. Fluorescence signals resulting from 470 nm and 415 nm excitation, interleaved in time, were sampled at 16 Hz.

For CSDS fiber experiments, videos were taken using Bonsai and handscored for attacks. For SIT and footshock, fiber signal was time-locked with video-tracking system (Ethovision XT 11, Noldus or ANY-Maze) via transistor-transistor logic signals (TTLs). Analysis was performed as previously described^1^.

**Behavior**

**Chronic Social Defeat Stress (CSDS) and Social Interaction Test (SIT)**

Mice were introduced into the cage of resident CD1 retired breeder mice pre-screened for aggression for 5 min/day over 10 days^2^ (Holt: 1,2). Following defeat, mice were placed on the other side of a perforated Plexiglass divider for the remaining 24 hours to allow sensory exposure. Control mice were housed with a Plexiglass divider between another control mouse (C57) and rotated to a different cage daily.

The SIT for social avoidance was performed within 24 hours after the last CSDS session. This test consisted of 2 portions run under red light conditions and animal exploratory behavior was assessed using Ethovision. First, mouse were placed in an empty open field arena containing a wire enclosure; the enclosure was empty for 2.5 mins (no target present). For the second portion, a novel CD1 mouse (non-aggressive) was placed in the wire enclosure (target present). Social interaction ratio was calculated by dividing the time spent in the interaction zone when the target mouse was present divided by the time spent in the zone when the target mouse was absent. Mice with a SI ratio < 0.9 were susceptible, and with > 1.1 were resilient^2,3^.

**Astrocyte Isolation**

Brains were extracted, sectioned into 1-mm-thick coronal slices using a brain matrix, and PFC tissue micropunched (unilateral 12G punches, Scientific Commodities; BB829-2-14). Astrocytes were immediately isolated using Miltenyi BioTech’s ACSA-2 MicroBead Kit (130-097-678) as previously described ^4-7^. Following astrocyte elution, a final 300 rcf centrifuge spin was performed, the supernatant removed, and astrocytes resuspended in 300 µL Tri-Reagent (Zymo R2050). To ensure lysis prior to snap-freezing on dry ice, the resuspension was vortexed for 30 seconds. All samples were stored at -80°C until RNA extraction.

RNA extraction, Library preparation, and Sequencing

RNA was extracted using Zymo’s Directzol RNA MicroPrep kit (R2061) following manufacturer’s instructions. RNA quality and quantity were assessed using Bioanalyzer (Agilent). Any sample with a RIN value of less than 7.8 was excluded. 500 pg of RNA was used as input in Takara’s SMARTer® Stranded Total RNA-Seq Kit v2 – Pico, with ribodepletion, according to the manufacturer’s instructions. Sequencing libraries were generated for each sample individually using Takara’s Unique Dual Index Kit. Following library preparation, sequencing was performed by Azenta on an Illumina Novaseq with a 2x150 bp paired-end read configuration to produce 40M reads per sample. Quality control was performed using FastQC.

**RNA extraction, library preparation, and sequencing**

RNA was extracted and prepped for sequencing as previously reported^2^. Differential gene expression (DEG) was performed in R (version 2025.05.1+513) using the DESeq2 package. Significant DEGs were determined by 20% log2foldchange and p < 0.05. Volcano plots, heatmaps, Venn diagrams, and gene ontology plots were generated using tidyverse and ggplot. Rank rank hypergeometric overlap (RRHO) were created using RRHO2 package. Gene ontology was performed using enrichR package. Top 10 terms were graphed presented and summarized for selected GO terms. Full GO terms are in supplemental table. For comparisons with human MDD and PTSD data sets, data was taken from Girgenti, et al.^8^ Supplemental Table. For correlations with behavior, variance-stabilized counts (VST) were obtained from DESeq2 analysis. For each gene, **Pearson correlation coefficient** was computed between its VST-normalized expression values and the corresponding behavioral metric. Correlations were calculated using pairwise complete observations to handle missing values.
