## Supplemental Figures for "Cortical astrocytes control stress resilience"

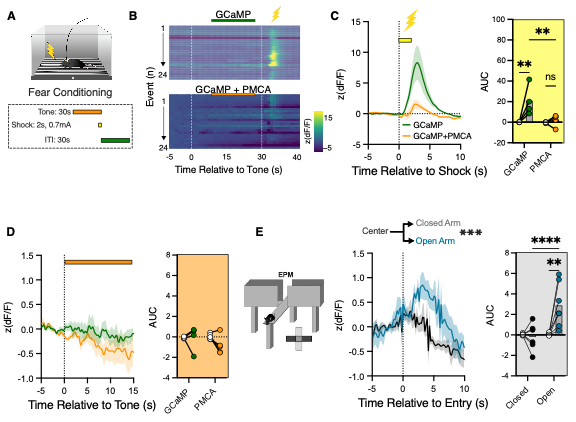

**Figure S1: PFC astrocytes respond to stressful and anxiogenic stimuli.**

A) Protocol for fear conditioning during fiber photometry recordings.
B) Heatmap of PFC astrocyte responses from GCaMP only and GCaMP-PMCA co-injected mice.
C) Average astrocytic response to footshock.
D) Resulting AUC to footshock.
E) Average astrocytic response to the shock-paired tone.
F) Resulting AUC.
G) Schematic of elevated plus maze (EPM).
H) Change in astrocytic Ca^2+^ when mice enter the closed arm vs. the open arm.
I) Resulting AUC upon entry to either arm of EPM.

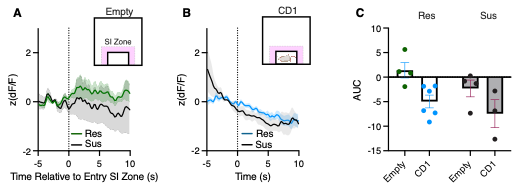

Figure S2: Astrocyte activity decreases during SIT, regardless of behavioral phenotypic profiling.

A) Difference in attack-evoked calcium activity in what will be susceptible verse resilient mice.
B) Change in astrocyte activity during interaction zone entry when chamber is empty.
C) Change in astrocyte activity when mice enter the interaction zone when a CD1 mouse is present.
D) Resulting AUC when mice enter social interaction zone.

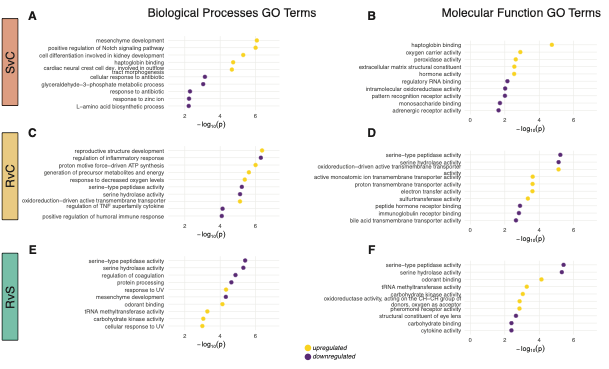

**Figure S3: Top 10 GO terms of comparisons.**

A) Biological processes GO terms comparing susceptible vs control mice.
B) Molecular function GO terms comparing susceptible vs control mice.
C) Biological processes GO terms comparing resilient vs control mice.
D) Molecular function GO terms comparing resilient vs control mice.
E) Biological processes GO terms comparing resilient vs susceptible mice.
F) Molecular function GO terms comparing resilient vs susceptible mice.

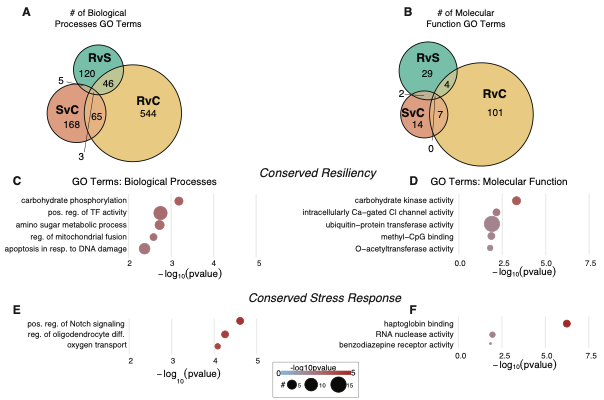

**Figure S4: Comparison of shared DEGs across behavioral phenotypic classification**

A) Overlap in biological processes GO terms across comparisons.
B) Overlap in molecular function GO terms across comparisons.
C) Biological process GO terms from resilient vs. control and resilient vs. susceptible comparisons, deemed conserved resilient genes.
D) Molecular function GO terms of conserved resilient genes.
E) Biological processes GO terms in shared DEGs between resilient vs. control and susceptible vs. control.
F) Molecular function GO terms in shared stress-responsive genes.

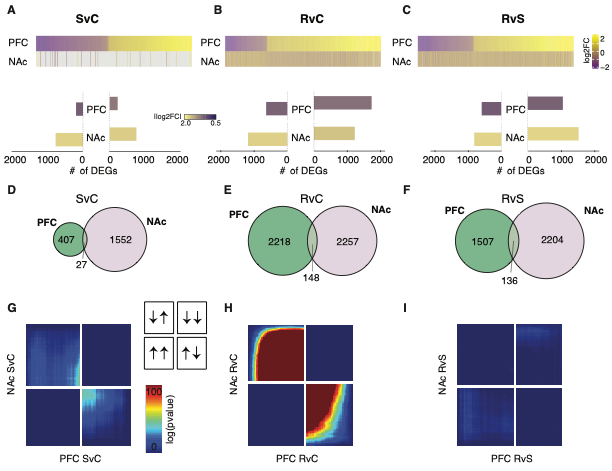

**Figure S5: Resilience induces distinct transcriptomic changes in astrocytes across limbic brain regions.**

A) Heatmap of DEGs from susceptible vs. control mice in PFC vs. NAc (top); plot of # of DEGs (bottom).
B) Heatmap of DEGs from resilient vs. control mice in PFC vs. NAc (top); plot of # of DEGs (bottom).
C) Heatmap of DEGs from resilient vs. susceptible mice in PFC vs. NAc (top); plot of # of DEGs (bottom).
D) Number of DEGs in PFC vs. NAc in susceptible vs. control mice.
E) Number of DEGs in PFC vs. NAc in resilient vs. control mice.
F) Number of DEGs in PFC vs. NAc in resilient vs. susceptible mice.
G) RRHO2 plot demonstrating patterns of gene changes comparing NAc vs. PFC susceptible vs. control mice.
H) RRHO2 plot of gene expression patterns in resilient vs. control mice comparing NAc vs. PFC.
I) RRHO2 plot of gene expression changes in PFC vs. NAc comparing resilient to susceptible mice.

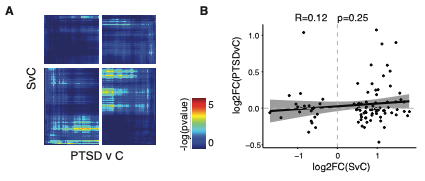

**Figure S6: PTSD in humans does not induce convergence with PFC astrocytes from susceptible mice.**

A) RRHO2 plot comparing PTSD with susceptible (vs control) mice.
B) Correlation of significant DEGs in susceptible vs control mouse conditions compared to PTSD in humans.
